# Catching SARS-CoV-2 by sequence hybridization: a comparative analysis

**DOI:** 10.1101/2021.02.05.429917

**Authors:** Alexandra Rehn, Peter Braun, Mandy Knüpfer, Roman Wölfel, Markus H. Antwerpen, Mathias C. Walter

## Abstract

Controlling and monitoring the still ongoing SARS-CoV-2 pandemic regarding geographical distributions, evolution and emergence of new mutations of the SARS-CoV-2 virus is only possible due to continuous next-generation sequencing (NGS) and worldwide sequence data sharing. Efficient sequencing strategies enabling the retrieval of the maximum number of high quality, full-length genomes are hence indispensable. Here, we describe for the first time a combined approach of digital droplet PCR (ddPCR) and NGS to evaluate five commercially available sequence capture panels targeting SARS-CoV-2. In doing so, we were not only able to determine the most sensitive and specific capture panel, but to discriminate their mode of action and number of read pairs needed to recover a high quality full length genome. Thereby, we are providing essential information for all sequencing laboratories worldwide striving for maximizing the sequencing output and simultaneously minimizing time, costs and sequencing resources.

## Introduction

At the moment, the world is still facing a tremendous and ongoing pandemic caused by a virus named SARS-CoV-2. While the mere detection by RT-qPCR or antigen tests to confine the spreading of this virus are valuable diagnostic tools, next-generation sequencing (NGS) techniques were, are and will be one of the keys to monitor and hence control this pandemic. Without the early availability of the SARS-CoV-2 genome in January 2020^1–3^ (strain Wuhan-Hu-1), the development of specific diagnostic RT-qPCR tests for the rapid detection of this virus would have been all but impossible^4^. At present, next-generation sequencing plus sharing the sequence data via the GISAID initiative is the only way to monitor the geographical distribution of the circulating strains and the adaption of the virus regarding its transmissibility^5–9^, pathogenicity^10–12^ and evolution^13,14^. Moreover, as antiviral treatments and vaccines have been developed against SARS-CoV-2, it is vital to know whether a newly emerged strain will develop resistance^15,16^ against antivirals or vaccine escaping mutations^17–19^.

However, direct NGS of human swab samples from COVID-19 positive patients, can be very expensive, time-consuming and challenging. Due to the fact that swab samples predominantly contain human cells with only a minor proportion of virus particles, direct sequencing of patient material is prone to missing the low-abundance species especially if no target enrichment strategies were applied prior to sequencing. At the moment, two different target enrichment approaches^20,21^ are mainly used around the world: Tiling multiplex PCRs^22–26^ and sequence hybridization by bait capture^27–29^. While the amplicon based enrichment is a very fast, sensitive and easy to handle approach, it can lead to sequencing gaps in case of divergences between the target genome and the amplicon primers due to mutations of the virus and is hence inconsistent in the elucidation of new SARS-CoV-2 mutations. Targeted capture-based approaches on the other hand tolerate up to 10-20% of mismatches between the target sequence and the so-called bait, which is made of biotinylated, single-stranded RNA/DNA probes complementary to the target DNA. Regarding the emergence of new SARS-CoV-2 mutations, we therefore see more certainty in using targeted-capture approaches. However, no evaluation of the various, commercially available capture bait panels has been conducted so far. We therefore set out to compare five different baits (Illumina Respiratory Panel v1 and v2, MyBaits SARS-CoV-2 Panel, Twist Bioscience SARS-CoV-2 Panel and Respiratory Panel) within three library preparation protocols in order to determine the most sensitive and most specific one, thereby providing pivotal information for all sequencing laboratories in the world that are currently occupied with the SARS-CoV-2 sequencing and the monitoring of new emerging mutations.

## Results

### Experimental Setup

In order to determine the sensitivity and specificity of the different baits (Illumina Respiratory Panel v1 and v2, MyBaits SARS-CoV-2 Panel, Twist Bioscience SARS-CoV-2 Panel and Respiratory Panel), five RNA input pools, varying in the ratio of the concentrations of SARS-CoV-2 and human reference RNA (HRR) to simulate human RNA background in patient samples, were produced. Absolute concentrations of SARS-CoV-2 and human RNA were quantified by droplet digital PCR (ddPCR) using the targets ORF1a and human ubiquitin C (UBC), respectively. The ORF1a to UBC ratio adjusted to 10^−5^ in pool 1, 10^−4^ in pool 2, 10^−3^ in pool 3, 10^−2^ in pool 4 and 10^−1^ in pool 5. The ratio of the produced input pools and the logarithmic change of the SARS-CoV-2 concentration were confirmed by ddPCR and reverse transcription-quantitative PCR (RT-qPCR) (see Suppl. Figure 1a, b). Subsequently, all RNA input pools were subject to reverse transcription and second strand synthesis before entering three different library preparation protocols provided by the companies Illumina, New England Biolabs (NEB) and Twist Bioscience (see Figure 1). Each library preparation was followed by an enrichment with a separate capture panel. In case of the Illumina library prep, the Respiratory Panels v1 and v2 from Illumina were used for the enrichment. The NEBNext Library preparation was followed by the sequence hybridization with the MyBaits SARS-CoV-2 panel, while the Twist Bioscience library preparation preceded the capture with the SARS-CoV-2 specific and the Respiratory Panel from Twist Bioscience (Figure 1). The change in the ratio of SARS-CoV-2 to human background was quantified by ddPCR before and after the capture. Finally, all enriched pools were sequenced on an Illumina MiSeq instrument.

**Figure 1:**
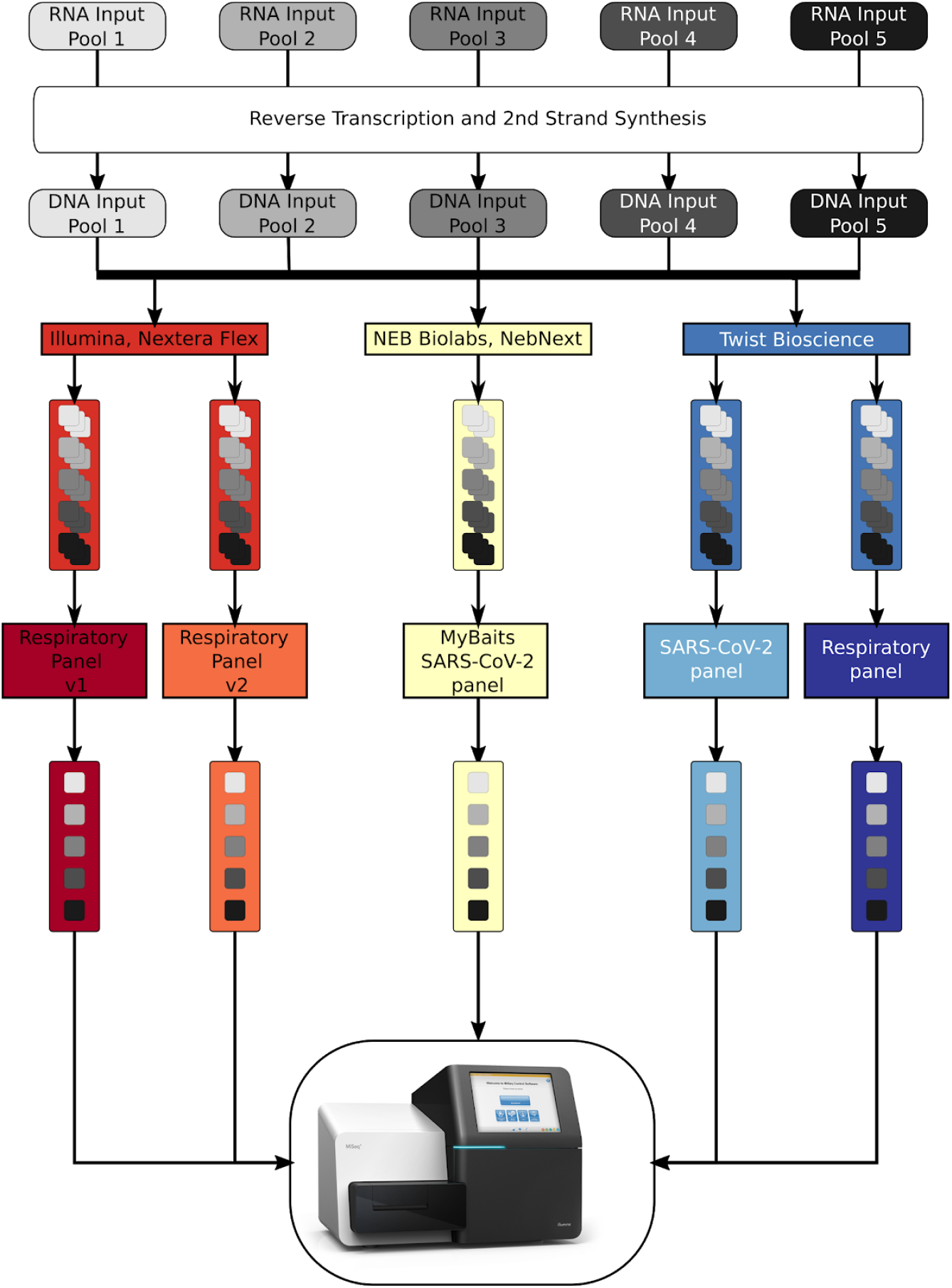
Graphical overview of the performed workflow. RNA input pools, varying in the SARS-CoV-2 concentration, were subject to reverse transcription and second strand analysis before entering three different library preparation methods. For better statistics, each RNA input pool was used three times during each library preparation method and for each bait panel tested. Before sequence hybridization, the triplicates were pooled by mass, resulting in 5 pools per bait, which were sequenced on an Illumina MiSeq after the enrichment process.

### Library preparation protocols differ significantly in quality and quantity of the processed library

Examination of the quality and quantity of the libraries is crucial for the subsequent sequencing and in this case for the consecutive target enrichment. Figure 2 shows the comparison of the libraries generated by the three different protocols with regard to fragment size, library concentration and total library mass. In terms of the mean fragment size and distribution, the Illumina Nextera Flex protocol produced the longest fragment with a mean length of about 600 bp, yet yielded the most atypical distribution as a second peak was visible in all samples (see Suppl. Figure 2). The libraries generated by the NEBNext and the Twist Bioscience protocols resulted in a mean fragment size of around 400 bp and 500 bp, respectively and showed a typical, Gaussian size distribution (Figure 2a and Suppl. Figure 2). Of note, all methods produced comparable fragment sizes across the pools 1-5, indicating highly reproducible procedures with a given input concentration. In contrast, the concentrations and thus the final library masses varied strongly between the three protocols (see Figure 2b, c). Here, the library preparation method of Twist Bioscience achieved the highest library concentrations surpassing their competitors by factor 1.4 and 7.5, respectively. Again, discrepancies between the pools were within the error range and indicate a stable and reproducible library preparation procedure for a given initial concentration. As single libraries ought to be pooled by mass prior to the sequence hybridization process according to the manufacturers’ protocols, a comparison of the final masses is beneficial (Figure 2c). Due to the highest library concentrations and the second highest elution volume, the library preparation method provided by Twist Bioscience resulted in the highest final library masses available for the subsequent sequence hybridization capture.

**Figure 2:**
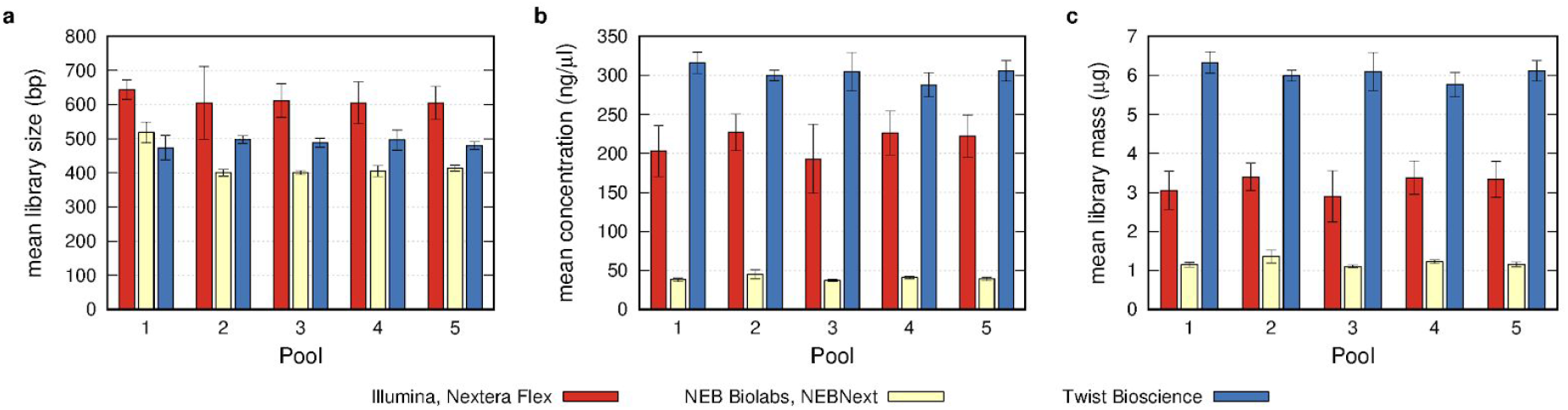
Comparison of the quality control parameters after library preparation with three different methods. a. Mean library size obtained by the analysis of the fragment size of the triplicates per input pool. Usage of the Illumina Nextera Flex protocol results in the largest libraries, followed by the libraries of Twist Bioscience and NebNext. b. Concentrations of the individual libraries were analysed with a Qubit fluorometer. Combining the values of the triplicates per input pool resulted in a mean concentration per pool. Here, the libraries produced by the Twist Bioscience protocol reached the highest mean concentration, followed by the libraries of the Illumina Nextera Flex and the NebNext protocol. c. Mean library mass was determined by the measured concentration and the elution volume. Here again, the Twist Bioscience libraries succeeded those of the Illumina Nextera Flex and NebNext

### Capture bait panels differ in their affinity towards SARS-CoV-2

In order to evaluate the sensitivity and specificity of the five different baits, the triplicates originating from the same RNA input pools were pooled by mass and quantified by ddPCR before and after the sequence hybridization process (Figure 3a and b). Primers targeting the ORF1a were used to quantify the presence of SARS-CoV-2 specific library fragments, while UBC was used as a marker for human non-target libraries. Pre-enrichment ORF1a:UBC ratios, depicted in Figure 3a reflect the exponential differences between the pools. Interestingly, the ORF1a:UBC ratio differed between the library preparation protocols, with the Illumina Nextera Flex protocol yielding the highest and the NEBNext libraries the lowest ORF1a:UBC ratio. The nature of this effect remains so far elusive and was not further addressed. Figure 3b shows the post-enrichment ORF1a:UBC ratio. Again, both Illumina panels showed the highest ORF1a:UBC ratio in all five pools, followed by the Twist Bioscience SARS-CoV-2 panel, the Twist Bioscience Respiratory Panel and the MyBaits SARS-CoV-2 panel. Moreover, all baits still reflected the exponential gradation of the ORF1a:UBC ratios from one pool to the next. To further discriminate the mode of action of the different bait panels during the enrichment process, the change of ORF1a- and UBC-concentration before and after the catch was compared using ddPCR (Figure 3c, d). Here, the Illumina bait panels achieved an ORF1a and hence a SARS-CoV-2 enrichment of about 100-fold. This together with the strongest depletion of UBC (Figure 3d), resulted in the highest ORF1a:UBC ratios after enrichment. Both panels from Twist Bioscience on the other hand yielded the strongest enrichment of ORF1a (Figure 3c) but were not able to decrease the UBC concentrations by more than one order of magnitude, especially the respiratory panel (Figure 3d). The MyBaits SARS-CoV-2 panel was neither efficient in the enrichment of ORF1a specific sequences nor in the depletion of UBC.

**Figure 3:**
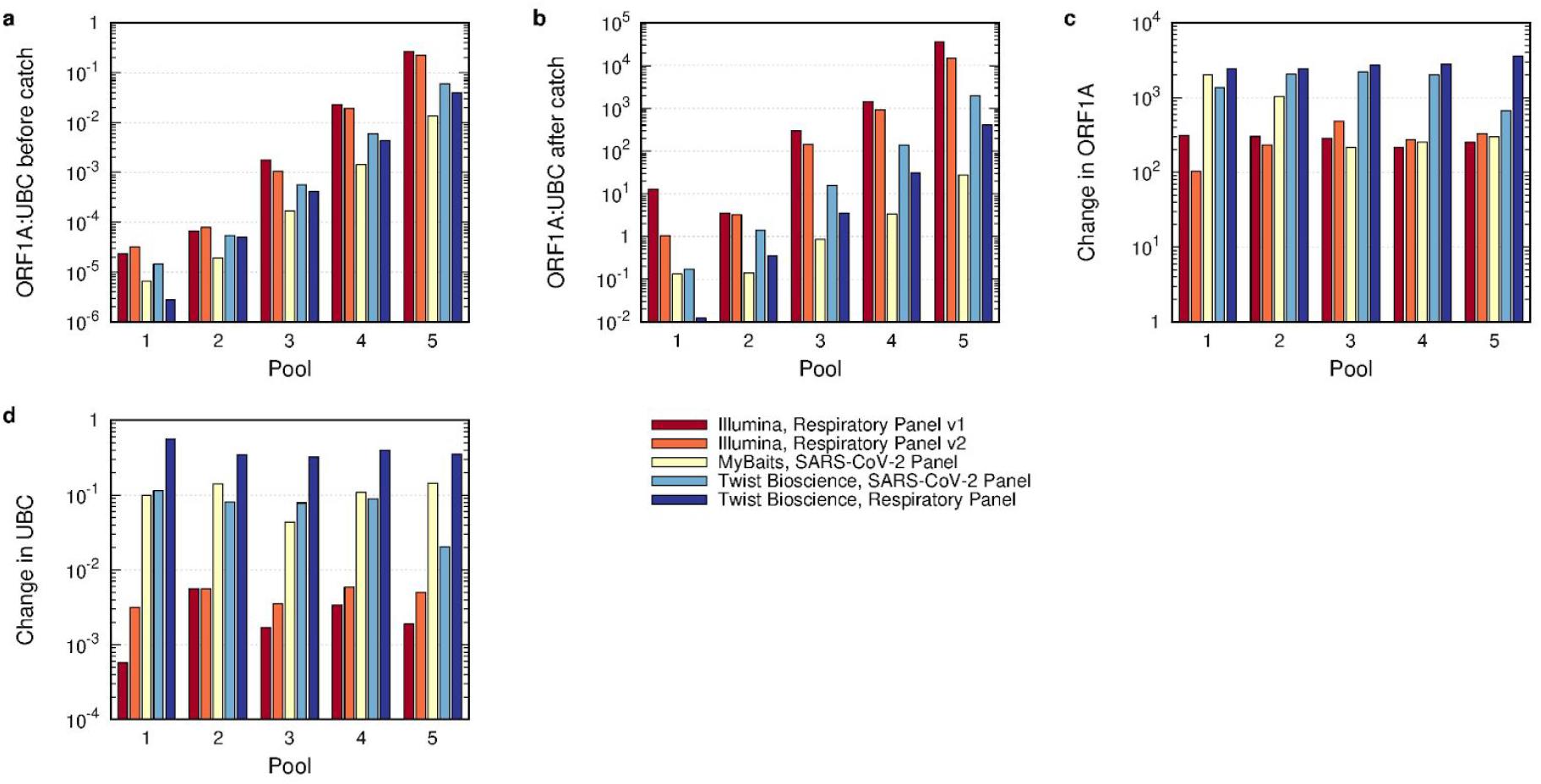
Analysis of the hybridization sequence capture by ddPCR. a and b: SARS-CoV-2 specific libraries were quantified by primers targeting ORF1A, while non-target libraries were quantified by the presence of human Ubiquitin C (UBC). The ORF1A:UBC ratio was plotted before (a) and after (b) the enrichment, showing the highest ratio for both Illumina panels, followed by both Twist Bioscience panels and the MyBaits panel. c and d: The change in ORF1A and UBC was plotted by dividing the counted concentration of ORF1A and UBC respectively after the enrichment with the respective concentrations before the enrichment. The strongest change in ORF1A was observed by both Twist Bioscience panels, while the strongest reduction of UBC was detected for the Illumina panels.

### Comparison of the sequence capture efficiency

After sequence hybridization, all enriched libraries were checked for concentration and fragment size (Suppl. Figure 3) and were subsequently sequenced on an Illumina MiSeq instrument. For an accurate comparison of the five different bait panels, all existing MiSeq reads were subsampled to 130,000 reads, which were previously corrected for PCR duplicates. All SARS-CoV-2 mapping reads within that subset were identified and the SARS-CoV-2:non-target ratio for each pool was plotted (Figure 4a). Usage of the SARS-CoV-2 specific bait panel of Twist Bioscience resulted in the highest abundance of SARS-CoV-2 specific reads in each pool, followed by the Respiratory Panel of Twist Bioscience and the Respiratory Panel v2 of Illumina, while the Respiratory Panel v1 and the MyBaits SARS-CoV-2 Panel produced the lowest number of SARS-CoV-2 specific reads (Figure 4a). Consistently, when applying the baits of the SARS-CoV-2 panel by Twist Bioscience nearly every base was already covered in pool 2 and resulted in a high-quality SARS-CoV-2 genome, in which every nucleotide of the genome was at least covered 20-fold, in pool 3 (Figure 4b and c). This was one pool and hence one order of magnitude earlier than the Respiratory Panel of Twist Bioscience and the Respiratory Panel v2 of Illumina, which were themselves another order of magnitude better than the Respiratory Panel v1 of Illumina and the MyBaits SARS-CoV-2 Panel (Figure 4b, c). In order to analyze the minimum number of reads needed to retrieve a full-length SARS-CoV-2 genome with a coverage of at least 20-fold (Table 1), the median coverage of the SARS-CoV-2 genome using 130,000 reads was calculated (see Suppl. Figure 4). In case the median coverage using 130,000 reads was zero, the median coverage of the SARS-CoV-2 genome was recalculated using all sequenced reads (indicated by a superscripted * in Table 1). If the median coverage was again zero, no number of reads could be assessed (indicated by N/A in Table 1). Table 1 shows that the usage of the SARS-CoV-2 panel from Twist Bioscience resulted in the least number of reads needed to recover a full-length SARS-CoV-2 genome. In fact, the number of reads was an order of magnitude lower than those of the Respiratory Panels from Twist Bioscience and Illumina, respectively, within the same pool. Again, the Respiratory Panel v1 and the MyBaits SARS-CoV-2 panel performed significantly less efficient than those previously mentioned.

**Figure 4:**
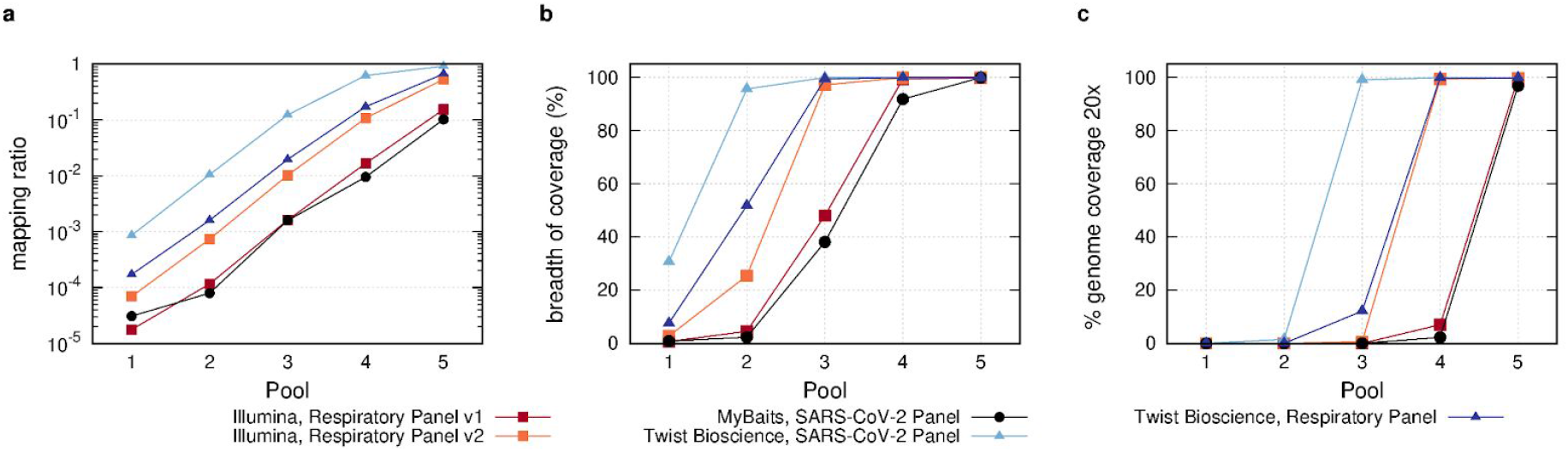
Analysis of the efficiency of the sequence hybridization panels by NGS. a: Number of SARS-CoV-2 mapping reads out of a subset of 130,000 reads were plotted against the pools, showing the highest mapping ratio for the Twist Bioscience SARS-CoV-2 panel. b: Breadth of coverage, defined by the number of covered bases of the SARS-CoV-2 genome, was compared for all panels. The usage of the Twist Bioscience SARS-CoV-2 panel led to a nearly complete coverage of the SARS-CoV-2 genome already in pool 2, while the Respiratory Panels of Twist Bioscience and Illumina reached the full breadth of coverage in pool 3. c: Comparison of the panels in regard to reaching a full length genome with a coverage of 20.

**Table 1.**
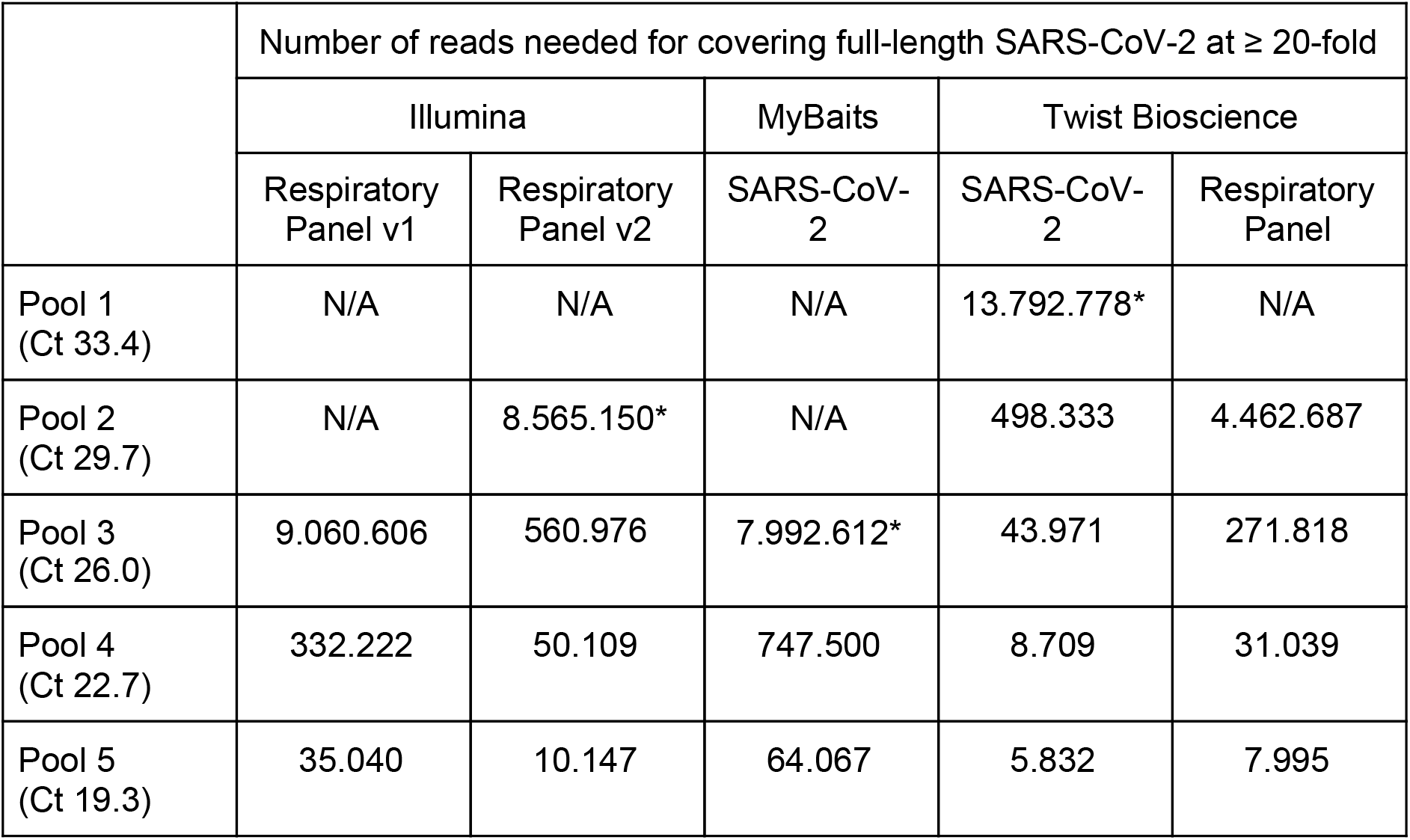
Overview of number of reads needed for retrieving a full length SARS-CoV-2 genome with a coverage of at least 20-fold regarding the choice of enrichment panel and the Ct values of the input pools. * Number of reads was calculated using the median coverage of all mapped reads instead of the subsampled mapped reads.

### Reasons for low capture rates and high non-target ratio

To further evaluate the source of the high number of non-target reads, especially in pools with a low input concentration of SARS-CoV-2, all non-target reads of all pools within a specific capture panel were mapped. Suppl. Table 1 shows a list of the top 30 hits of all panels, thereby revealing mainly ribosomal RNA (rRNA) targets when sorting by the number of total hits. These reads account for the majority of panels between 56% and 97% of all non-target reads, with the exception of the Respiratory Panel v1 of Illumina in which only 7.3% hits are caused by rRNA (Table 2). Interestingly, the highest number of hits (about 25%) in this panel was assigned to GAPDH, which was drastically reduced in the successor version v2 and is obsolete in the capture panels of the other companies. Nevertheless, this analysis reveals a room for improvement either in the capture panels themselves or in the complete strategy by adding a rRNA depletion step.

**Table 2.** Overview of number of non-target reads and their major hits

|  | Illumina |  | Twist Bioscience |  | MyBaits |
| --- | --- | --- | --- | --- | --- |
| Panel | Respiratory Panel v1 | Respiratory Panel v2 | SARS-CoV-2 | Respiratory Panel | SARS-CoV-2 |
| non-target reads | 2,527,450 | 1,896,363 | 1,286,780 | 2,186,252 | 4,817,333 |
| rRNA reads | 183,994 | 1,066,572 | 811,090 | 2,132,833 | 3,010,428 |
| rRNA : non-target | 7.3% | 56.2% | 63.0% | 97.6% | 94.1% |
| GAPDH reads | 630,000 | 176,046 | 0 | 0 | 0 |
| GAPDH : non-target | 24.9% | 9.3% |  |  |  |

## Discussion

The SARS-CoV-2 pandemic, which originated in Wuhan in December 2019 is still ongoing and reached, despite the use of numerous counteractive measures, new records of infected individuals in December 2020. Since the beginning of the pandemic, whole genome sequence data generated by next-generation sequencing were shared publicly on platforms like GISAID and played a pivotal role in the identification^1^, the development of diagnostic^4^ and therapeutic^30,31^ strategies, the investigation of the origin^13^ and the evolution of the virus. Driven by the appearance of potentially more aggressive, more infectious/contagious or immunity escaping strains like B1.1.7 (UK)^6,32^, B1.315 (South Africa)^7,18^ and P1 (alias of B.1.1.28.1, Brazil)^33,34^, the world health organization (WHO) initiated in January 2021 a sequencing program^35^ to monitor the viral movement, activity and evolution with its impact on transmissibility, pathogenicity and immunity. In order to reach these goals, a large number of SARS-CoV-2 genomes will need to be sequenced continuously and efficiently in terms of time and costs. Therefore, target enrichment protocols like capture-based or amplicon-based approaches are inevitable and allow more samples to be sequenced in parallel^36^. While the amplicon-based enrichment (ARTIC^23^ und Co.), which generates target amplicons from 400-2,000 bp, is very sensitive, it is also more prone to amplicon failure due to divergences in the target genome at the primer binding sites, leading to gaps in the genome sequence and hence loss of potentially important information, especially when looking for new mutations. Targeted capture-based approaches on the other hand are able to tolerate up to 10-20% of mismatches between the target sequence and the bait^35^, thereby providing a stable technique in the monitoring of new SARS-CoV-2 variants. We therefore set out to compare five different capture baits towards their sensitivity and specificity to SARS-CoV-2 by a combined approach of ddPCR and NGS.

Our results demonstrate that all tested baits were able to bind SARS-CoV-2 libraries but showed great differences in their enrichment capacities. Altogether, the SARS-CoV-2 Panel of Twist Bioscience performed best followed by the Respiratory Panel from Twist Bioscience,the Respiratory Panel v2 from Illumina, its progenitor panel v1 and the MyBaits SARS-CoV-2 panel. We speculate that this hierarchy is a result of the combination of three parameters: First, the enrichment factor for ORF1a/SARS-CoV-2 reads, secondly the depletion factor for non-target reads and last but not least the fragment size after the sequence hybridization. The SARS-CoV-2 specific panel from Twist Bioscience showed together with the Respiratory Panel from Twist Bioscience the highest enrichment factor for ORF1a/SARS-CoV-2, but succeeded the Respiratory Panel in the depletion of UBC/non-target reads (Figure 3c,d). Additionally, all Twist Bioscience libraries displayed the largest post-enrichment fragment size (Suppl. Figure 3b), thereby rendering both panels as the best and second-best performing ones. The Illumina Respiratory Panel v1 and v2 on the other hand, showed only an enrichment factor for ORF1a/SARS-CoV-2 of about 100 fold, but performed best in the depletion of the UBC/non-target reads (Figure 3c,d). Nevertheless, the post-enrichment fragment size of the Illumina libraries was significantly smaller than the one from Twist Bioscience (Suppl. Figure 3b). NGS data of the Respiratory Panel v2 showed a higher number of target reads (Figure 4a), thereby surpassing the older version v1. The MyBaits SARS-CoV-2 panel was the only capture-based approach sold as a stand-alone product without any recommended library preparation protocol. Here, we observed that the combination of NEBNext Ultra II library preparation protocol and the MyBaits SARS-CoV-2 panel resulted in the least sensitive combination with the lowest specificity. Our data clearly revealed that the NEBNext library prep resulted in the shortest libraries with the lowest concentrations. Whether this was the main cause for the poor performance or the combination of the baits with this library preparation is impossible to tell from our data, since the combination of the best performing Twist Bioscience SARS-CoV-2 baits with the NEBNext libraries was not performed.

This is to date the first study comparing capture enrichment panels for SARS-CoV-2. We were able to identify the best performing one and also successfully deconstructed the mode of SARS-CoV-2 enrichment and depletion of non-target reads between the different panels. By combining this information we are proposing on the one hand an improvement of the capture efficiency by either adding a rRNA depletion step or by removing individual bait sequences that are responsible for the targeting of rRNA molecules. On the other hand, our study provides a correlation between the SARS-CoV-2 concentration (measured by RT-qPCR or ddPCR) and the minimal number of reads needed to recover a high-quality full-length genome, thereby reducing valuable time, sequencing resources and costs. Hence, this work may pave the way for high throughput yet high quality screening for the worldwide emerging new mutations of SARS-CoV-2 and hence contribute to a more effective containment of the ongoing Covid19 pandemic.

## Methods

### Cultivation and Purification of SARS-CoV-2

SARS-CoV-2 virus was cultured in Vero E6 cells with MEM containing 2% FBS at 37° C with 5% CO_2_ and was harvested 72 hours post infection. Virus stocks were stored at −80° C. Viral RNA was extracted using diatomaceous earth^37^. Briefly, 140 μL of virus-containing supernatant was added to 560 μL lysis buffer (800 mM Guanidine hydrochloride, 50 mM Tris [pH 8.0], 0.5% Triton X-100, 1% Tween-20) and incubated at room temperature for 10 min. Subsequently, 560 μL ethanol (VWR) as well as 20 μL diatomaceous earth (VWR, 100 mg/mL in distilled water) were added to the mixture. After vigorous vortexing, the diatomaceous earth cell culture mixture was incubated at room temperature for 5 min with shaking to prevent sedimentation of the diatomaceous earth. After centrifugation at 13,000 rpm for 3 min at room temperature, the supernatant was discarded. 500 μL of washing buffer (10 mM Tris [pH 8.0], 0.1% Tween-20) was added, and the mixture was centrifuged at 13,000 rpm for 3 min. After discarding the supernatant, 500 μL of washing buffer (10 mM Tris [pH 8.0], 0.1% Tween-20) was added, and the mixture was centrifuged again for 3 min at 13,000 rpm. After decanting the supernatant, 400 μL of acetone (Roth) was added to the pellet, vortexed and centrifuged again. After removing the supernatant, the pellet was dried for 5 min at 56° C and the viral RNA was eluted with 80 μL of distilled water. After mixing and centrifugation, the RNA was transferred to a new reaction tube and stored at −80° C until further use.

### Quantification of SARS-CoV-2 and human RNA and cDNA by droplet digital PCR (ddPCR) and reverse transcription (RT) ddPCR

For quantification of human Ubiquitin C mRNA (UBC) and SARS-CoV-2 Open Reading Frame 1a (ORF1a) RNA, 20 μl ddPCR mix consisted of 5μl One-Step RT-ddPCR Advanced Supermix for Probes (Bio-Rad, Laboratories, Munich, Germany), 2 μl of Reverse Transcriptase (Bio-Rad, final concentration 20 U/μl), 1 μl of DTT (Bio-Rad, Laboratories, Munich, Germany; final concentration 15 nM), 1 μl 20x UBC primer and probe mix (Table 3, final concentrations: primers 900 nM, probe 250 nM), 1 μl of 20x ORF1a primer and probe mix (Table 3, final concentrations: primers 900 nM, probe 250 nM), 5 μl of nuclease free water (Qiagen, Hilden, Germany) and 5 μl of template RNA. Partitioning of the reaction mixture into up to 20,000 droplets was carried out on a QX200 ddPCR droplet generator (Bio-Rad, Laboratories, Munich, Germany) and PCR was performed using a Mastercycler Pro (Eppendorf, Wesseling-Berzdorf, Germany) with the following thermal protocol: Reverse transcription was performed at 50° C for 60 min. An enzyme activation step at 95° C was carried out for 10 min followed by 40 cycles of a two-step program of denaturation at 94° C for 30 s and annealing/extension at 58° C for 1 min. Final enzyme inactivation was performed at 98° C for 10 min. Finally, the samples were cooled down to 4°C. All steps were performed using a temperature ramp rate of 2° C/s. After PCR droplets were analyzed using a QX100 Droplet Reader (Bio-Rad, Laboratories, Munich, Germany) and Quantasoft Pro Software was used (Bio-Rad, Laboratories, Munich, Germany) for absolute quantification of target concentrations.

**Table 3.**
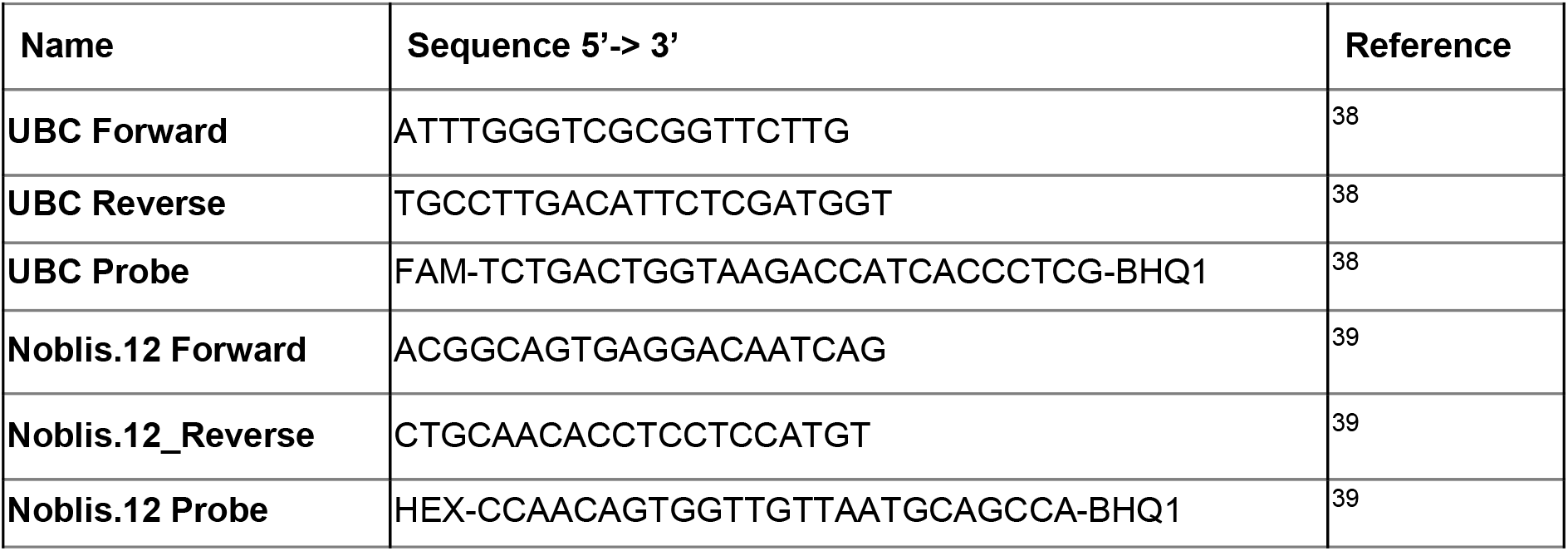
Primers and probes used in this study.

When cDNA was used as a template the 20 μl ddPCR mix consisted of 10 μl ddPCR Supermix for Probes (Bio-Rad Laboratories, Munich, Germany), 1 μl 20x UBC primer and probe mix (Table 3, final concentrations: primers 900 nM, probe 250 nM), 1 μl of 20x ORF1a primer and probe mix (Table 3, final concentrations: primers 900 nM, probe 250 nM), 3 μl of nuclease free water (Qiagen, Hilden, Germany) and 5 μl of template containing cDNA. Subsequent steps were carried out as described for RT-ddPCR with the difference that no initial reverse transcription step was included in the thermal cycling protocol.

### Generation of RNA Input Pools

In order to create RNA pools with varying SARS-CoV-2 concentrations, the initial concentrations of purified SARS-CoV-2 and the universal human reference RNA (UHRR, Agilent Technologies, product number 740000) were determined by ddPCR as described above. Subsequently, each RNA input pool was calculated to have a SARS-CoV-2 to UBC ratio of 10^−5^ in pool 1, 10^−4^ in pool 2, 10^−3^ in pool 3, 10^−2^ in pool 4 and 10^−1^ in pool 5. Evaluation of the SARS-CoV-2:UBC ratio of these RNA input pools was again done by ddPCR.

### Reverse Transcription and Second Strand Synthesis

Dependent on the subsequent library preparation protocol, two different reverse transcriptases were used. In the case of Illumina Nextera Flex and NEB Biolabs NEBNext, SuperScript IV (ThermoFisher) was applied according to the manufacturers’ recommendations, while ProtoScript II (NEB Biolabs) was used for the Twist Bioscience workflow according to the details given in the Twist Bioscience Library Preparation protocol. To improve the efficiency of all reverse transcriptases, the random hexamers (Random Primer 6 in case of Protoscript II) were mixed with 10% Oligo(dT) primers. In all cases, the NEBNext Ultra II Non-directional RNA Second Strand Synthesis Buffer and Reagents (NEB Biolabs) was used for the second strand synthesis.

### Library Preparation

Library preparation was performed according to the manufacturers’ protocols. For Illumina, the Nextera Flex for Enrichment (Version v03) was used with the following deviations: In the step “Amplify Tagmented DNA”, the initial denaturation time was prolonged from 3 min to 4 min. Furthermore, the denaturation time during the 12 cycles of amplification was set to 30 sec instead of 20 sec. For the preparation of the Twist Bioscience libraries the guide “Creating cDNA Libraries using Twist Library Preparation Kit for ssRNA Virus Detection” (version: August 2020) was followed according to the instructions given. In step 3.1, the fragmentation time was reduced from 22 min to 1 min. For NebNext libraries, the manual “NEBNext Ultra II FS DNA Library Prep Kit for Illumina” was used. Here, we followed the instructions of section 1 for inputs ≤ 100 ng and also reduced the fragmentation time to 1 min.

### Sequence Capture by Hybridization

In order to compare the five bait panels, 200 ng of each triplicate were pooled and subsequently hybridized according to the manufacturer’s instruction. In case of the Respiratory Bait Panels v1/v2 from Illumina, the hybridization was performed at 58° C and overnight. After washing, the enriched libraries were amplified for 12 cycles. Here, the initial denaturation time was prolonged to 60 sec, while the denaturation time during the cycles was set to 20 sec. For enrichment of the Twist libraries with either the SARS-CoV-2 specific or the Respiratory Panel, the manual “Twist Target Enrichment Protocol” was followed without any exception. Similarly, the MyBaits “Hybridization Capture for Targeted NGS” manual (version 4.01) was used according to the manufacturer’s instructions to enrich the NebNext libraries.

### Quality Control of Libraries and Sequencing

After library preparation and after the enrichment, the libraries had to pass a quality control check regarding concentration and size. The concentrations of the libraries were measured on a Qubit 4 fluorometer using the ds DNA HS Assay Kit (Thermo Fisher Scientific). The shape and the mean fragment size of the libraries were determined on a 5200 Fragment Analyzer using the HS NGS Fragment Kit 1-6000 bp (both Agilent). Enriched libraries were loaded with a final concentration of 10 pM on a MiSeq Flow Cell using v3 reagent chemistry for 2×150 cycles.

### Data Analysis

Sequenced reads were cleaned from PCR duplicates using clumpify from the BBTools package^40^ prior subsampling them to 130,000 reads using seqtk^41^ to get normalized datasets for each pool. Afterwards, subsampled reads were mapped against the SARS-CoV-2 Wuhan-Hu-1 reference genome sequence^1^ with GenBank accession MN908947.3 using bwa mem^42^. The number of mapped reads were determined using samtools flagstat^43^ and coverage information was obtained using `bedtools genomecov`^44^. Data collection and overall statistics were generated using custom bash and awk scripts. Datamash^45^ was used to aggregate the triplicate datasets and gnuplot^46^ for plotting.

To get a near-optimal pool ratio in correlation to the library concentration, we estimate the number of reads needed for covering a full-length SARS-CoV-2 genome at a minimum of 20-fold by simply solving the triangle inequality of 130,000 reads divided by the median sequence coverage of the pool and multiplied with the target coverage of 20. The result was further corrected by the number of observed PCR duplicates.

To investigate the high number of reads not mapping to the SARS-CoV-2 genome a combined FASTA file containing all human reference genome sequences, the SARS-CoV-2 reference genome sequence as well as all annotated RNAs (non-coding and mRNAs) of both genomes was created. Then, Salmon^47^ was used with default settings (kmer=31) to quantify the transcript abundance of all sequenced reads of each triplicate against this dataset. Transcripts targeted by more than 100 reads were extracted and aggregated for each pool. For the 30 top-most targeted transcripts their gene name and function were looked up and further aggregated if transcripts belong to the same gene.

## Notes

### Competing Interest Statement

The authors have declared no competing interest.

### Summary of Updates

Typographic and grammar corrections

