## Supplementary Data for "Catching SARS-CoV-2 by sequence hybridization: a comparative analysis" (from another version of this preprint: version 1, 2021-02-05)

### Suppl. Fig. 1: ddPCR Input Pools, qPCR Input Pools

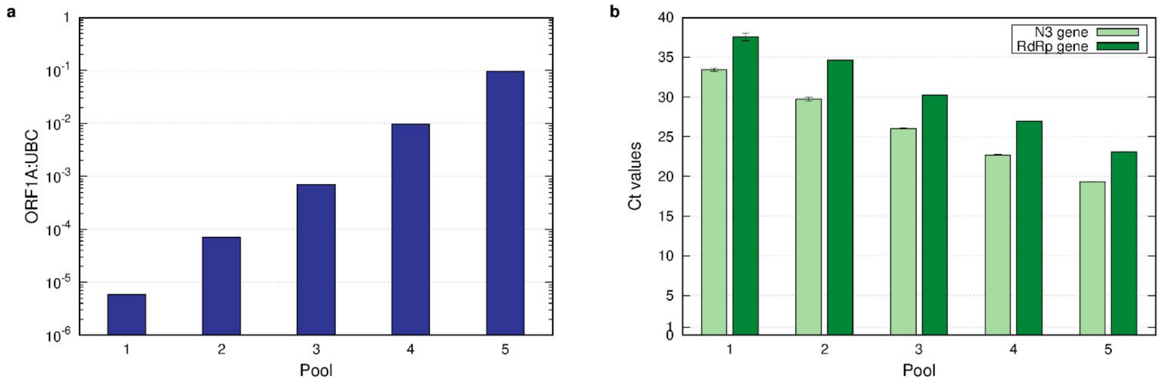


### Suppl. Fig. 2: Fragment Analyzer of Illumina, NebNext, Twist

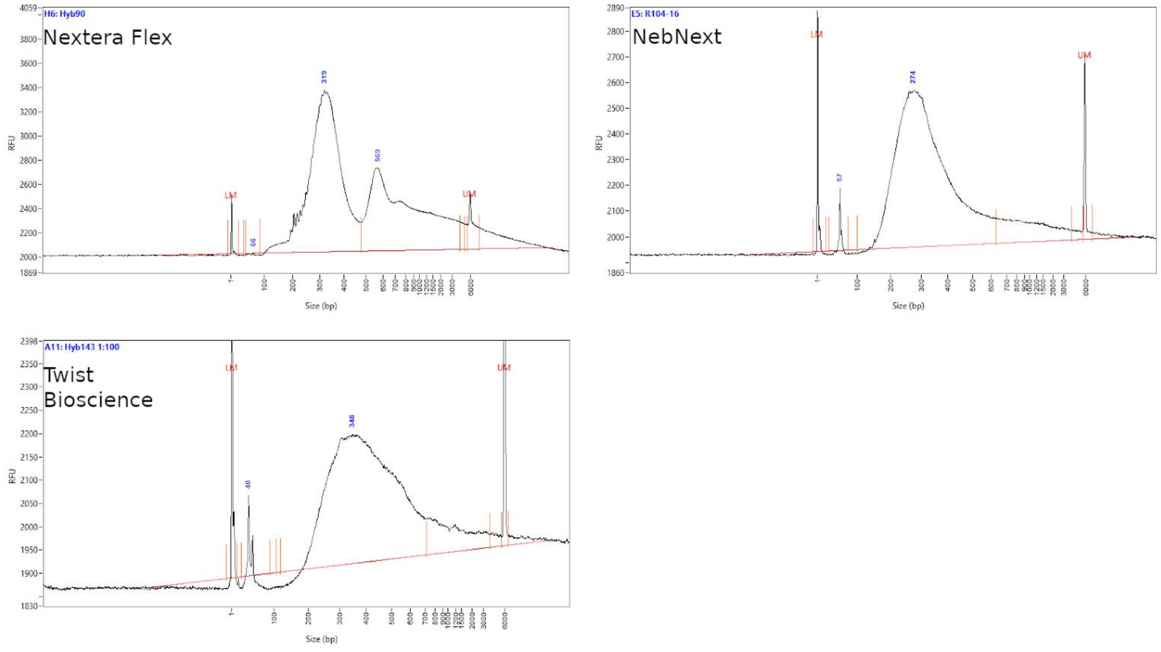


### Suppl. Fig. 3: QC der enriched libraries

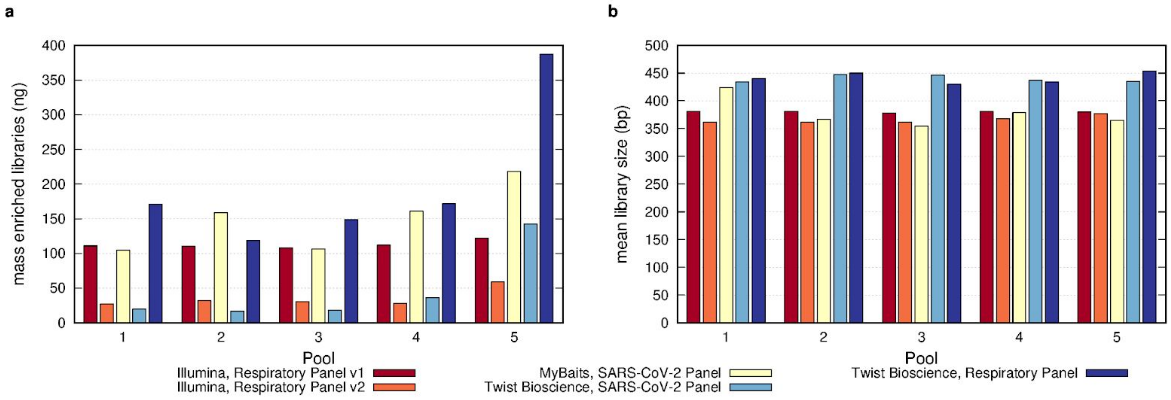

Suppl. Fig. 4:

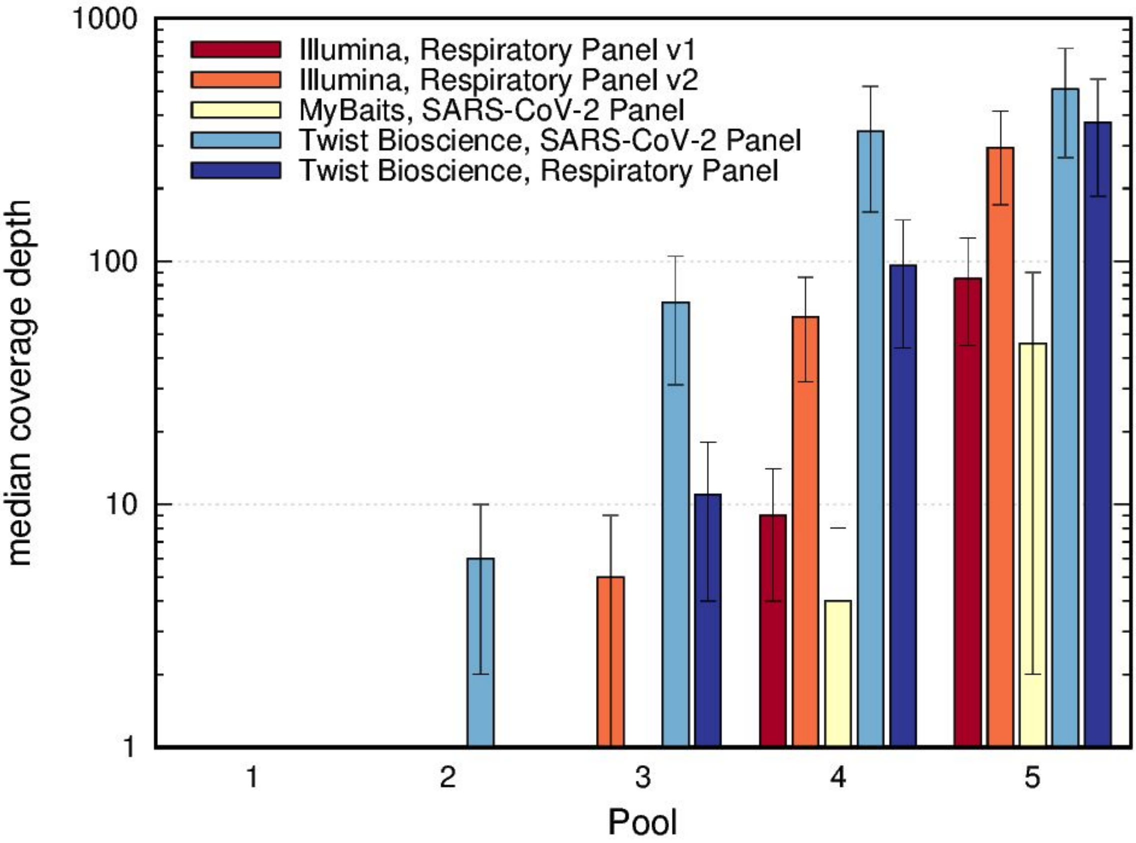

Illumina, Respiratory Panel v1
Illumina, Respiratory Panel v2
MyBaits, SARS-CoV-2 Panel
Twist Bioscience, SARS-CoV-2 Panel
Twist Bioscience, Respiratory Panel
median coverage depth
1000
100
10
1
1
2
3
4
5
Pool

**Suppl. Table 1.** The 30 top-most captured transcripts are shown with their aggregated number of reads per enrichment panel and pool.

| | | | | Illumina | | Twist Bioscience | | MyBaits | |
|---|---|---|---|---|---|---|---|---|---|
| Accession | Type | Gene | Product | Respiratory Panel v1 | Respiratory Panel v2 | SARS-CoV-2 | Respiratory Panel | SARS-CoV-2 | Total reads / transcript |
| NR_146118.1 | rRNA | RNA28SN4 | RNA, 28S ribosomal N4 | 11,580 | 124,670 | 267,055 | 687,946 | 1,015,800 | 2,107,051 |
| NR_146119.1 | rRNA | RNA18SN4 | RNA, 18S pre-ribosomal N4 | 10,091 | 96,756 | 202,794 | 541,354 | 843,533 | 1,694,528 |
| NR_145822.1 | rRNA | RNA28SN1 | RNA, 28S ribosomal N1 | 28,121 | 107,255 | 205,439 | 587,204 | 751,128 | 1,679,147 |
| NR_152150.2 | ncRNA | GAPDH | glyceraldehyde-3-phosphate dehydrogenase | 516,290 | 151,119 | | | | 667,410 |
| NR_146117.1 | rRNA | RNA45SN4 | RNA, 45S pre-ribosomal N4 | 95,232 | 471,221 | 5,958 | 34,831 | 44,836 | 652,078 |
| NR_146148.1 | rRNA | RNA28SN2 | RNA, 28S ribosomal N2 | | 8,702 | 61,154 | 124,193 | 230,357 | 424,406 |
| NM_002796.3 | mRNA | PSMB4 | proteasome 20S subunit beta 4 | 164,428 | 48,499 | | | | 212,927 |
| NR_003287.4 | rRNA | RNA28SN5 | RNA, 28S ribosomal N5 | | | 52,286 | 75,762 | 57,608 | 185,657 |
| NR_145819.1 | rRNA | RNA45SN1 | RNA, 45S pre-ribosomal N1 | 6,271 | 136,769 | 4,205 | 11,869 | 16,335 | 175,449 |
| NM_001289745.3 | mRNA | GAPDH | glyceraldehyde-3-phosphate dehydrogenase | 113,709 | 24,927 | | | | 138,636 |
| NR_146151.1 | rRNA | RNA45SN3 | RNA, 45S pre-ribosomal N3 | 30,724 | 98,201 | | 1,153 | 5,780 | 135,858 |
| NR_146154.1 | rRNA | RNA28SN3 | RNA, 28S ribosomal N3 | 1,064 | 4,308 | 6,846 | 37,262 | 13,099 | 62,578 |
| NM_004637.6 | mRNA | RAB7A | RAB7A, member RAS oncogene family | 77,349 | 16,996 | | | | 94,345 |
| NM_002787.5 | mRNA | PSMA2 | proteasome 20S subunit alpha 2 | 77,041 | 17,273 | | | | 94,314 |
| NM_002793.4 | mRNA | PSMB1 | proteasome 20S subunit beta 1 | 73,953 | 20,251 | | | | 94,204 |
| NR_146144.1 | rRNA | RNA45SN2 | RNA, 45S pre-ribosomal N2 | 910 | 18,690 | 5,354 | 31,259 | 31,952 | 88,165 |
| XM_017028118.1 | mRNA | EIF2S2 | eukaryotic translation initiation factor 2 subunit beta | 71,881 | 12,815 | | | | 84,696 |
| NM_015388.4 | mRNA | YIPF3 | Yip1 domain family member 3 | 58,909 | 15,660 | | | | 74,569 |
| NM_006827.6 | mRNA | TMED10 | transmembrane p24 trafficking protein 10 | 59,292 | 13,109 | | | | 72,401 |
| NM_015383.2 | mRNA | NBPF14 | neuroblastoma breakpoint family (NBPF) member 14 | | | 68,241 | 4,133 | | 72,374 |
| NM_005482.3 | mRNA | PIGK | phosphatidylinositol glycan anchor biosynthesis class K | 11,583 | 60,354 | | | | 71,937 |
| XR_002959730.1 | miscRNA | LRPAP1 | LDL receptor related protein associated protein 1 | 56,604 | 12,484 | | | | 69,088 |
| NM_032847.3 | mRNA | C8orf76 | chromosome 8 open reading frame 76 | 9,916 | 52,661 | | | | 62,577 |
| NM_032015.5 | mRNA | RNF26 | ring finger protein 26 | 44,012 | 13,306 | | | | 57,318 |

| | | | | | | | | | |
|---|---|---|---|---|---|---|---|---|---|
| NM_001042559.3 | mRNA | EIF4G2 | eukaryotic translation initiation factor 4 gamma 2 | 44,027 | 7,828 | | | | 51,855 |
| NM_002806.5 | mRNA | PSMC6 | proteasome 26S subunit, ATPase 6 | 36,495 | 9,619 | | | | 46,114 |
| NM_015084.3 | mRNA | MRPS27 | mitochondrial ribosomal protein S27 | 36,086 | 9,767 | | | | 45,853 |
| NM_005702.4 | mRNA | ERAL1 | Era like 12S mitochondrial rRNA chaperone 1 | 34,612 | 8,438 | | | | 43,050 |
| NM_032940.3 | mRNA | POLR2C | RNA polymerase II subunit C | 33,466 | 9,210 | | | | 42,676 |
| NM_178819.4 | mRNA | GPAT4 | glycerol-3-phosphate acyltransferase 4 | 28,122 | 9,584 | | | | 37,706 |
| NM_025147.4 | mRNA | COQ10B | coenzyme Q10B | 5,788 | 31,036 | | | | 36,825 |
